# SR Ca^2+^ handling in unbranched, immediately post-necrotic fast-twitch *mdx* fibres is similar to *wt* littermates

**DOI:** 10.1101/2021.11.29.470495

**Authors:** Stephen Chan, Sindy L. L. Kueh, John W. Morley, Stewart I. Head

**Affiliations:** School of Medicine, Western Sydney University, Campbelltown, New South Wales, Australia; Department of Physiology, Faculty of Science, Mahidol University, Ratchatewi, Bangkok, Thailand

**Keywords:** SR calcium, *mdx* mouse, branched fibres

## Abstract

There is a lack of consensus in the literature regarding the effects of dystrophin deficiency on the Ca^2+^-handling properties of the SR in *mdx* mice, an animal model of Duchenne muscular dystrophy. One possible reason for this is that only a few studies control for the presence of branched fibres. Fibre branching, a consequence of degenerative-regenerative processes such as muscular dystrophy, has in itself a significant influence on the function of the SR. In our present study we attempt to detect early effects of dystrophin deficiency on SR Ca^2+^ handling by using unbranched fibres from the immediate post-necrotic stage in *mdx* mice (just regenerated following massive necrosis). Using kinetically-corrected Fura-2 fluorescence signals measured during twitch and tetanus, we analysed the amplitude, rise time and decay time of Δ[Ca^2+^]_i_ in unfatigued and fatigued fibres. Decay was also resolved into SR pump and SR leak components. Fibres from *mdx* mice were similar in all respects to fibres from *wt* littermates apart from: (i) a longer rise time and slower rate of rise of [Ca^2+^]_i_ during a tetanus; and (ii) a mitigation of the fall in Δ[Ca^2+^]_i_ amplitude during the course of fatigue. Our findings suggest that the early effects of a loss of dystrophin on SR Ca^2+^ handling are only slight, and differ from the widely held view that there is significant Ca^2+^ pathology in *mdx* mice. It may be that Ca^2+^ pathology is magnified by progressive branching and degeneration.

**New findings:** Central question: What are the early effects of dystrophin deficiency on SR Ca^2+^ handling in the *mdx* mouse?

Main finding: In the *mdx* mouse, Ca^2+^ handling by the SR is little affected by the absence of dystrophin when looking at fibres without branches that have just regenerated following massive myonecrosis. This has important implications for the traditional view that Ca^2+^ pathology is significant in the *mdx* mouse.

## Introduction

Impairment of Ca^2+^ handling is recognised as a key event in the pathogenesis of Duchenne muscular dystrophy (DMD) (Burr & Molkentin, 2015). This degenerative muscle disease results from a complete lack of the long isoform of dystrophin, a rod-shaped protein from the inner surface of the sarcolemma of skeletal muscle fibres (Emery, 2003). In the *mdx* mouse, an animal model of DMD that also lacks the long isoform of dystrophin, abnormal Ca^2+^ fluxes have been observed across both the sarcolemma and the sarcoplasmic reticulum (SR) (reviewed in Vallejo-Illarramendi *et al.* 2014). Given its role as the Ca^2+^ regulator of skeletal muscle (Rossi & Dirksen, 2006), the SR is one organelle where signs of disrupted Ca^2+^ handling might be expected to manifest, even though it is spatially separated from dystrophin. Indeed, alterations of Ca^2+^ release, uptake, storage and leakage have been found in the SR of *mdx* mice.

A reduction of Ca^2+^ release in *mdx* mice has been reported by Woods *et al.* (2004) and Hollingworth *et al.* (2008), who detected a reduced amplitude of the Ca^2+^ transient and a reduced Ca^2+^ release flux, and by Plant & Lynch (2003), who found a reduced caffeine-induced calcium release in skinned fibres. Ca^2+^ uptake in skinned fibres of *mdx* mice has been reported to be slower (Divet & Huchet-Cadiou, 2002) or faster (Robin *et al.* 2012) than *wt*. Levels of the Ca^2+^ storage protein calsequestrin in *mdx* mice are reduced in the diaphragm but increased in muscles spared from degeneration, such as the extraocular muscles (Pertille *et al.* 2010). The SR of dystrophin-deficient fibres is thought to be especially leaky to Ca^2+^ following mechanical stress, exhibiting greater Ca^2+^ spark activity than control fibres following eccentric contraction (Bellinger *et al.* 2009) or osmotic shock (Wang *et al.* 2005). An increased SR Ca^2+^ leak has also been found in unstressed skinned fibres from *mdx* mice (Divet & Huchet-Cadiou 2002; Robin *et al.* 2012).

Other studies, however, have reported that some features of SR Ca^2+^ handling are not affected by dystrophin deficiency. Plant & Lynch (2003) found no change in the Ca^2+^ uptake and Ca^2+^ leak of the SR in skinned fibres from *mdx* mice. Collet *et al.* (1999) found that the twitch transients in fibres from *wt* and *mdx* mice were similar in all respects apart from the decay kinetics in response to a short-duration depolarising pulse.

One possible reason for this lack of consensus is that the muscle fibres in the *mdx* mouse are not homogeneous. One property that varies between muscle fibres is the degree of fibre branching. Branched fibres are a consequence of muscle regeneration (Schmalbruch, 1976). At any one time, a muscle from an *mdx* mouse contains some fibres that are branched and some that are not, and those that are branched have varying numbers and complexity of branches (Chan & Head, 2011).

At 3-4 weeks of age, *mdx* mouse hindlimb muscles undergo a massive necrosis and regeneration that replaces virtually all muscle fibres (Duddy *et al.* 2015). The necrotic-regenerative process stabilises to a low rate by 6-8 weeks of age (Grounds *et al.* 2008), at which time about 15% of fibres in extensor digitorum longus (EDL) and 5% of fibres in flexor digitorum brevis (FDB) are branched. Almost all branches are single. This stable rate of necrosis persists throughout life, with the proportion of branched fibres increasing to 90% in EDL and 30% in FDB after 6 months of age. The branches also become multiple and more complex (Head *et al.* 1992; Chan *et al.* 2007).

Branching in itself affects the SR Ca^2+^ handling of a muscle fibre. The action-potential-evoked Ca^2+^ transient has a lower amplitude in branched *mdx* muscle fibres than in unbranched *mdx* fibres (Head, 2010; Lovering *et al.* 2009; Hernández-Ochoa *et al.* 2015). Within a branched *mdx* fibre, the amplitude of the Ca^2+^ transient is lower in the branch portion than in the trunk portion (Lovering *et al.* 2009; Hernández-Ochoa *et al.* 2015). The decay time of the Ca^2+^ transient is longer in a branched *mdx* fibre than in an unbranched one (Lovering *et al.* 2009). During tetanic stimulation at high frequencies, *mdx* fibres fail to maintain a Ca^2+^ plateau, and this failure occurs at a lower stimulation frequency in branched than in unbranched fibres (Head, 2010). Ca^2+^ spark activity induced by osmotic stress is greater in the branch portion than in the trunk portion of a branched *mdx* fibre (Lovering *et al.* 2009).

It is therefore possible that the disagreement over the effects of dystrophin deficiency on SR Ca^2+^ handling is at least partly due to variations in the proportions of branched fibres in study samples. The muscle selected, the age of the mouse and the portion of the fibre that is imaged could all affect the likelihood of branches being included in the data. In order to more accurately compare dystrophin-deficient with dystrophin-positive fibres, it is therefore desirable to minimise the confounding effects of fibre branching on the Ca^2+^ signals measured.

To this end, we conduct our present study, which examines the SR Ca^2+^ handling of unbranched muscle fibres from *mdx* mice aged 7 weeks. This is the earliest age at which the rate of myonecrosis reaches its stable lifetime level. We thus attempt to minimise the effects of branching or other pathologies which are downstream of the initial dystrophin defect, in order to more clearly isolate the effect of dystrophin deficiency on SR Ca^2+^ handling.

Other SR Ca^2+^ studies have explicitly controlled for branched fibres (Lovering *et al.* 2009; Hernández-Ochoa *et al.* 2015), but we extend their analyses of unbranched fibres by: (i) using even younger mice; (ii) separating the decay of the Ca^2+^ transient into SR pump and SR leak components; (iii) examining tetanic stimulation; and (iv) examining fatiguing stimulation. In addition, we use wild-type littermates as the control animal for the *mdx* mice, a practice which, to our knowledge, has been followed only by Bellinger *et al.* (2009) in SR Ca^2+^ studies.

## Methods

### Animals used

Animals were used in accordance with Western Sydney University Animal Care and Ethics Committee approval A12346. In contrast with the vast majority of studies using *mdx* mice, we used *wt* littermates as our controls (Kueh *et al.* 2004; Kiriaev *et al.* 2009). Male dystrophic mice and littermate controls were obtained from the Western Sydney University animal facility. The colony used in this study were second-generation offspring of C57BL/10SnSn DMD (*mdx*) mice. Mice were genotyped to distinguish dystrophin-deficient from control mice (Amalfitano & Chamberlain, 1996). All animals used were aged 7 weeks.

### Tissue preparation

Animals were sacrificed with an overdose of halothane. The flexor digitorum brevis (FDB) muscle was dissected from the hindlimb and incubated in a muscle digest solution for 30 min at 37°C. The digest solution was a Krebs solution (4.75 mM KCl, 118 NaCl, 1.18 KH_2_PO_4_, 1.18 MgSO_4_, 24.8 NaHCO_3_, 2.5 CaCl_2_ and 10 glucose) to which was added 3 mg/mL collagenase I (Sigma Chemical Co., St Louis, MO, USA) and 1 mg/mL trypsin inhibitor (Sigma), aerated with 95% O_2_ - 5% CO_2_ to maintain pH at 7.4. Following incubation, the muscle mass was washed twice in Krebs-only solution. Individual fibres were then obtained by gently pipetting the muscle mass (Head *et al.* 1990).

The results shown here are for 7 fibres from 3 *wt* animals and 7 fibres from 4 *mdx* animals.

### Fluorescence measurements

#### Dye ionophoresis

The fibres were placed onto glass coverslips for fluorescence microscopy and became firmly attached. Individual muscle fibres were viewed with a 40 UV-F objective on a Nikon TE300 inverted microscope equipped a xenon light source. Fibres with diameter >40 μm were selected; these larger fibres were the MT-II fast Ca^2+^ transient type (Calderón *et al.* 2009). Among these fibres, only those that gave a quick, reproducible twitch response to electrical stimulation were used for experimentation. An intracellular electrode was used to fill the muscle fibres with the ionised form of the Ca^2+^-sensitive dye fura-2. Fura-2 (1 mM) in distilled H_2_O was introduced into the tip of the ionophoretic electrode, and the shank was then filled with 150 mM potassium acetate. Dye was ionophoresed into the muscle fibres to give a final concentration of 5–50 μM fura-2 in the cell (Head, 1993). After filling with fura-2, the fibres were left for about 20 min before any readings were taken to allow for complete distribution of the dye in the myoplasm.

#### Fluorescence sampling

Fluorescence emission intensities at 510 nm were sampled via a photo multiplier tube (PMT) at 20,000 Hz using a spectrophotometer (Cairn) under 360 and 380 nm excitation. Fibres were stimulated in three basic modes: (i) twitches; (ii) graded frequency run at 2, 5, 10, 15, 20, 25, 30 and 50 Hz stimulation frequency for 500 ms each; or (iii) fatigue run consisting of 50-Hz stimulation for 500 ms every 1.0 s for 60 s. In each mode, recordings were made at 380 nm first, and then repeated at 360 nm. The 360-nm wavelength was found to be very close to the isosbestic value and was used as an indicator of movement artifact. This was negligible for twitches but noticeable for some tetani; in any case the ratio of 360- to 380-nm signals was used in [Ca^2+^]_i_ calculations (see below).

#### Fibre stimulation

The fibre was stimulated using a bipolar stimulating electrode placed close to the neuromuscular junction, which was visible in the light microscope as a corrugated oval on the fibre. The fibre was stimulated with pulses of 1 ms duration from 1 to 50 Hz. Shortening in response to action potential activation of the fibres was minimal. During the experiments, the isolated fibres were superfused (1 mL/min) with Krebs solution maintained at room temperature (22°C – 24°C) and aerated with 95% O_2_ - 5% CO_2_.

#### [Ca^2+^]_i_ calculation

In this study, we were interested in the changes in [Ca^2+^]_i_ during stimulation and hence the ratio of 360- to 380-nm signals would have been sufficient to compare many calcium parameters of *wt* and *mdx*. However, in order to perform kinetic correction and SR pump function analysis (see below), it was necessary to generate [Ca^2+^]_i_ values. This was done using the equation of Grynkiewicz *et al.* (1985):

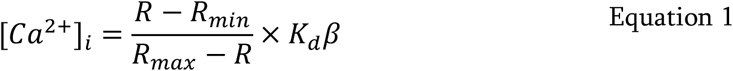

where *R* is the measured ratio of 360- to 380-nm fluorescence, *R_min_* is the ratio under Ca^2+^-free conditions, *R_max_* is the ratio at saturating [Ca^2+^], *K_d_* is the apparent dissociation constant of fura-2 and *β* is the ratio of the 380-nm fluorescence under minimum and maximum [Ca^2+^]. Values were assigned to *R_min_*, *R_max_* and *K_d_β* in order to give a resting [Ca^2+^]_i_ of 45 nM at the start of each experiment. This was reasonable based on previous measurements (Head, 1993) and provided values that could be used to compare changes in [Ca^2+^]_i_ between *wt* and *mdx* during stimulation.

#### Kinetic correction of fura-2 transients

Because of the slow binding kinetics of fura-2, very fast events such as Ca^2+^ release during muscle stimulation are not adequately captured, with a marked underestimation of the rate of Ca^2+^ release. This limitation can be overcome by applying a correction process to the calculated [Ca^2+^] values, as outlined in Bakker *et al.* (1997):

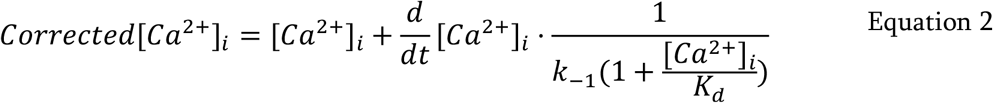

where *k*_-1_ is the dissociation constant of Ca^2+^-fura-2. We used this correction in calculating the kinetics of the rising phase of the Ca^2+^ transients. Following correction, the rise times of the Ca^2+^ transients obtained in this study are in good agreement with those obtained by Calderón *et al.* (2009) for type IIX fibres from mouse FDB muscle using faster, lower-affinity dyes.

### Statistics

Data are presented as Mean ± S.E.M.. Comparison of *wt* and *mdx* parameters before fatigue was conducted using two-tailed *t*-tests. Comparison of *wt* and *mdx* parameters after fatigue and after recovery was conducted using two-way ANOVA with Sidak’s correction for multiple comparisons. Comparison of *wt* and *mdx* parameters during fatiguing stimulation was conducted by fitting a least-squares regression curve to the changing values over time, and then comparing the curves. If a suitable equation could not be found for fitting the values, the comparison was made by two-way ANOVA with Sidak’s correction for multiple comparisons. All tests were conducted at a significance level of 5%. All statistical procedures were performed using a standard statistical software package (GraphPad Prism Version 8 for Windows, GraphPad Software, San Diego California USA).

## Results

### Experimental protocol

The experimental protocol is illustrated in Figure 1 using examples from both a *wt* littermate control and an *mdx* animal. A series of twitches (Stage A) was followed immediately by a graded frequency run (Stage B) in which the fibre was stimulated at 2, 5, 10, 15, 20, 25, 30, 50 and 100 Hz. Only the 50-Hz tetanus is shown in the figure. Then came the fatigue run (Stage C) consisting of 50-Hz stimulation for 0.5 s every 1.0 s until 60 s had elapsed. This was immediately followed by a couple of twitches (Stage D) and another graded frequency run (Stage E). After 3 min recovery (Stage F), there was another set of twitches (Stage G) and graded frequency run (Stage H). Throughout the text, these stages will be referred to as pre-fatigue (A and B), intra-fatigue (C), post-fatigue (D and E) and post-recovery (G and H).

**Figure 1.**
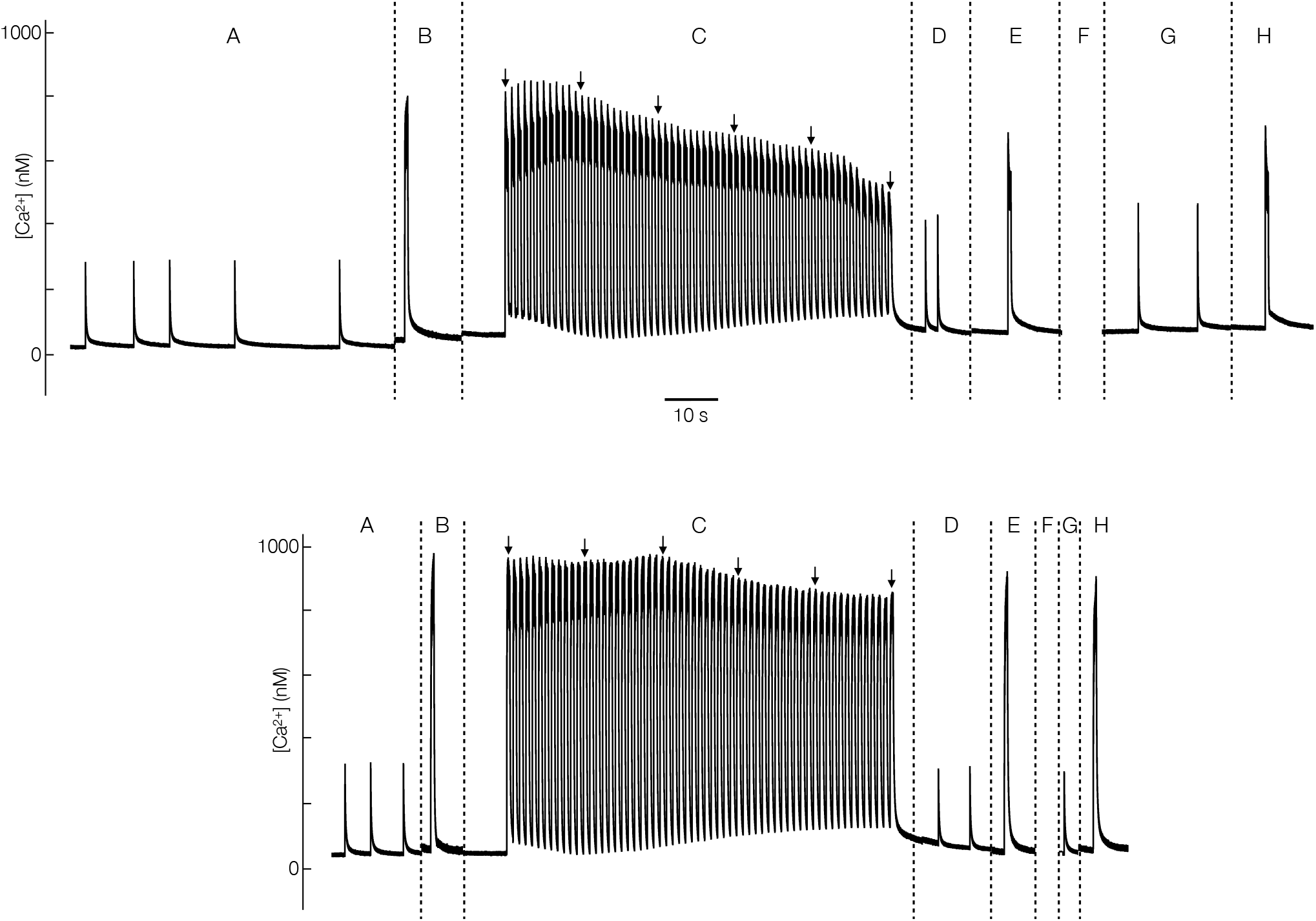
The experimental protocol. *Upper trace* from *wt* animal aged 7 wks. *Lower trace* from *mdx* animal aged 7 wks. (A) Series of twitches. (B) Graded frequency run with stimulation at 2, 5, 10, 15, 20, 25, 30 and 50 Hz. Only 50-Hz tetanus shown here. (C) Fatigue run with stimulation at 50 Hz for 500 ms, every 1 s for 60 s. (D) One or two twitches. (E) Graded frequency run. Only 50-Hz tetanus shown here. (F) Recovery for 3 mins. (G) Series of twitches. (H) Graded frequency run. Only 50-Hz tetanus shown here. *Arrows* point to tetani at 0, 12, 24, 36, 48 and 60 s into fatigue run.

In all fibres included in these results, the level of the post-recovery [Ca^2+^]_i_ approached or even exceeded the pre-fatigue levels. This was true even when the experimental protocol was repeated, up to five times in some cases.

### [Ca^2+^]_i_ changes during a twitch

The changes in [Ca^2+^]_i_ during a twitch are illustrated in Figure 2. If more than one twitch was recorded, the data were averaged to obtain a single tracing. Figure 2A shows an averaged fura-2 recording (*black line*) and the kinetically corrected [Ca^2+^]_i_ (*red line*) which has been adjusted for fura-2 binding kinetics. The kinetically corrected [Ca^2+^]_i_ shows a spike that is much larger and peaks earlier than the uncorrected [Ca^2+^]_i_, and a decay phase that follows the same course as the decay of the uncorrected [Ca^2+^]_i_. Thus to measure very fast events such as Ca^2+^ release, the kinetically corrected values must be used, but to measure slower events such as Ca^2+^ uptake, the uncorrected values can be used.

**Figure 2.**
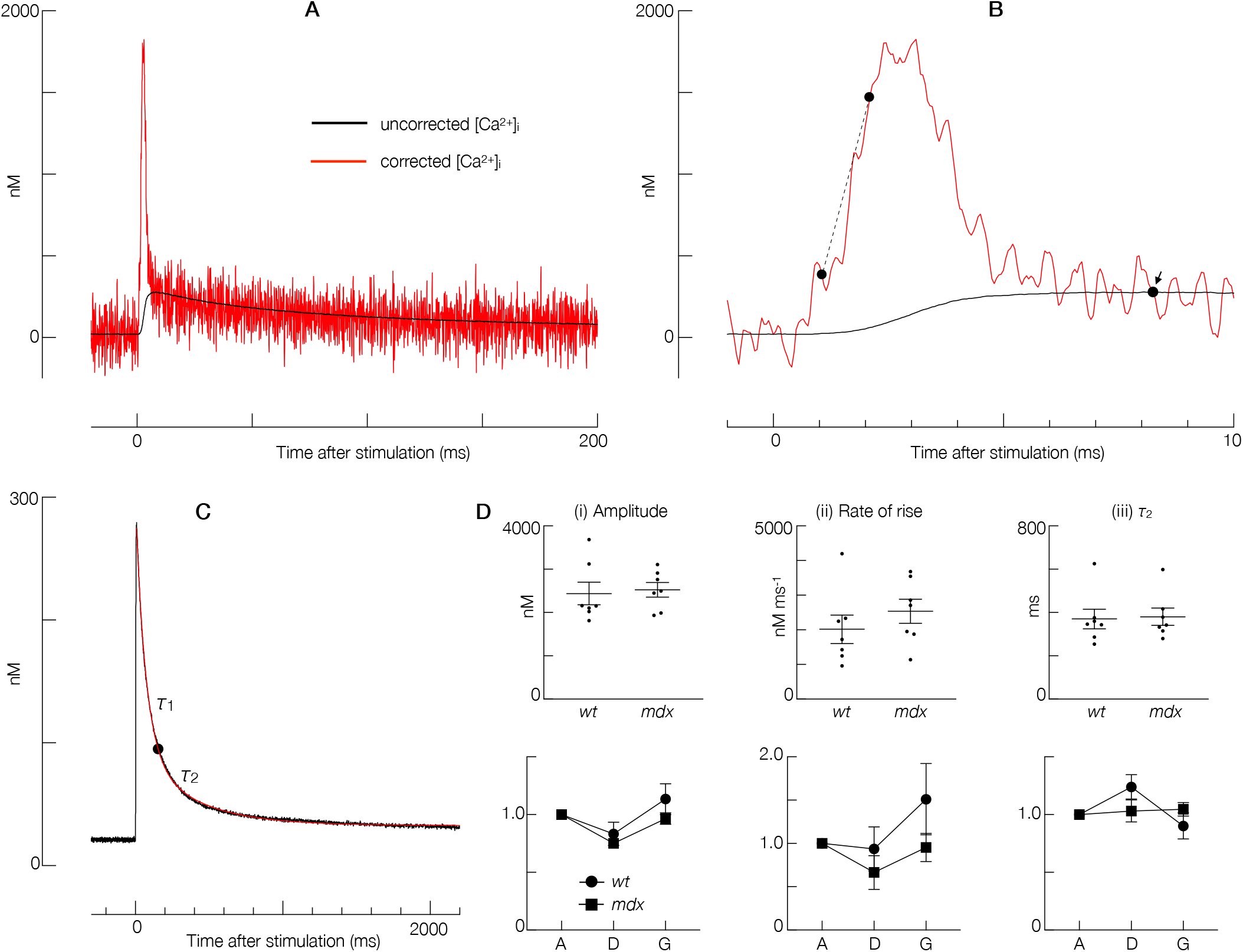
Changes in [Ca^2+^]_i_ during a twitch. **(A)** Effect of kinetic correction. *Black line* uncorrected [Ca^2+^]_i_. *Red line* kinetically corrected [Ca^2+^]_i_. It can be seen that the kinetically corrected [Ca^2+^]_i_ consists of a spike, followed by a decay that matches the decay of the uncorrected [Ca^2+^]_i_. **(B)** The first 10 ms of the data shown in (A), focusing on the spike of the corrected [Ca^2+^]_i_. *Dotted line* joins the points at which [Ca^2+^]_i_ reaches 20% and 80% of the [Ca^2+^]_i_ amplitude. *Arrow* indicates the peak of the uncorrected [Ca^2+^]_i_. **(C)** *Black line* same uncorrected [Ca^2+^]_i_ as in (A), but showing the full time course of decay. *Red line* fitted double exponential function with time constants *τ*1 and *τ*2. Data for (A), (B) and (C) are taken from the same experiment as in Figure 1. **(D)** Summary statistics for (i) amplitude of [Ca^2+^]_i_; (ii) rate of rise of [Ca^2+^]_i_ (slope of the dotted line in (B)); and (iii) *τ*_2_. *Upper row* pre-fatigue results (from Stage A — see Figure 1). *Lower row* effects of fatiguing stimulation and recovery. Labels on *x*-axis refer to the stages of the experiment described in Figure 1. Values on *y*-axis are relative values, with Stage A given a value of 1.0.

#### Spike amplitude

The statistics for the amplitude of the spike in the kinetically corrected [Ca^2+^]_i_ are shown in Figure 2Di. The *upper graph* shows pre-fatigue values; the *lower graph* shows post-fatigue and post-recovery values, relative to pre-fatigue values. No significant difference was found between *wt* littermate controls and *mdx* before fatigue. As a whole, the spike amplitude was reduced following the fatigue run and increased back up to pre-fatigue levels following 3 minutes of recovery. However, the extent of these changes was not different between *wt* and *mdx*.

#### Rate of rise of spike

Figure 2B is an expansion of the first 10 ms of Figure 2A and illustrates how the kinetically corrected [Ca^2+^]_i_ reaches its peak much earlier than the uncorrected [Ca^2+^]_i_. The rate at which the corrected [Ca^2+^]_i_ rises is measured by the slope of the dotted line, which joins the points at which [Ca^2+^]_i_ reaches 20% and 80% of its amplitude. Figure 2Dii shows that this rate of rise is not significantly different between *wt* littermate controls and *mdx* before fatigue, and the changes following fatigue and recovery are also similar between the two genotypes.

#### Slow time constant of decay

Figure 2C shows the whole decay phase of the same uncorrected [Ca^2+^]_i_ as in Figure 2A. A double exponential function was fitted (*red line*), yielding the time constant *τ*_2_ of the slow phase of decay. Figure 2Diii shows that there was no difference between *wt* littermate controls and *mdx* in pre-fatigue values of *τ*_2_, nor was there any difference in the changes resulting from fatigue and recovery.

### SR pump function

The SR pump function curve is an accepted methodology which separates the decay of [Ca^2+^]_i_ into two factors — SR Ca^2+^ pumping and SR Ca^2+^ leak (Klein *et al.* 1991, Allen & Westerblad 1995).

Figure 3A shows uncorrected [Ca^2+^]_i_ of twitches from three stages of the same experiment: pre-fatigue, post-fatigue and post-recovery. The recordings are shown only from the time when [Ca^2+^]_i_ falls below 150 nM. The slow phase of decay, determined as in Figure 2C, begins shortly after this time. To improve the fit even further, a double exponential fit was applied again, just to this slow phase, and the resulting curves are the *red lines*. At several arbitrary points along these red lines, the values of [Ca^2+^]_i_ and -*d*[Ca^2+^]_i_/*dt* were determined, and these values are plotted on Figure 3B. Equation 3 (Klein *et al.* 1991) was then fitted to these points, resulting in

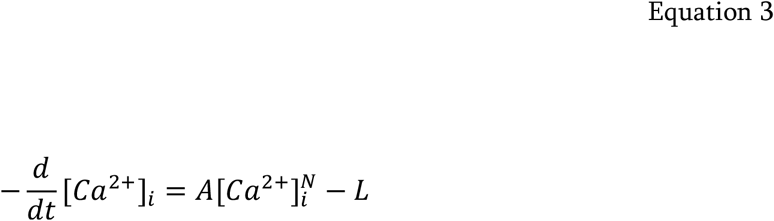

the curves shown.

**Figure 3.**
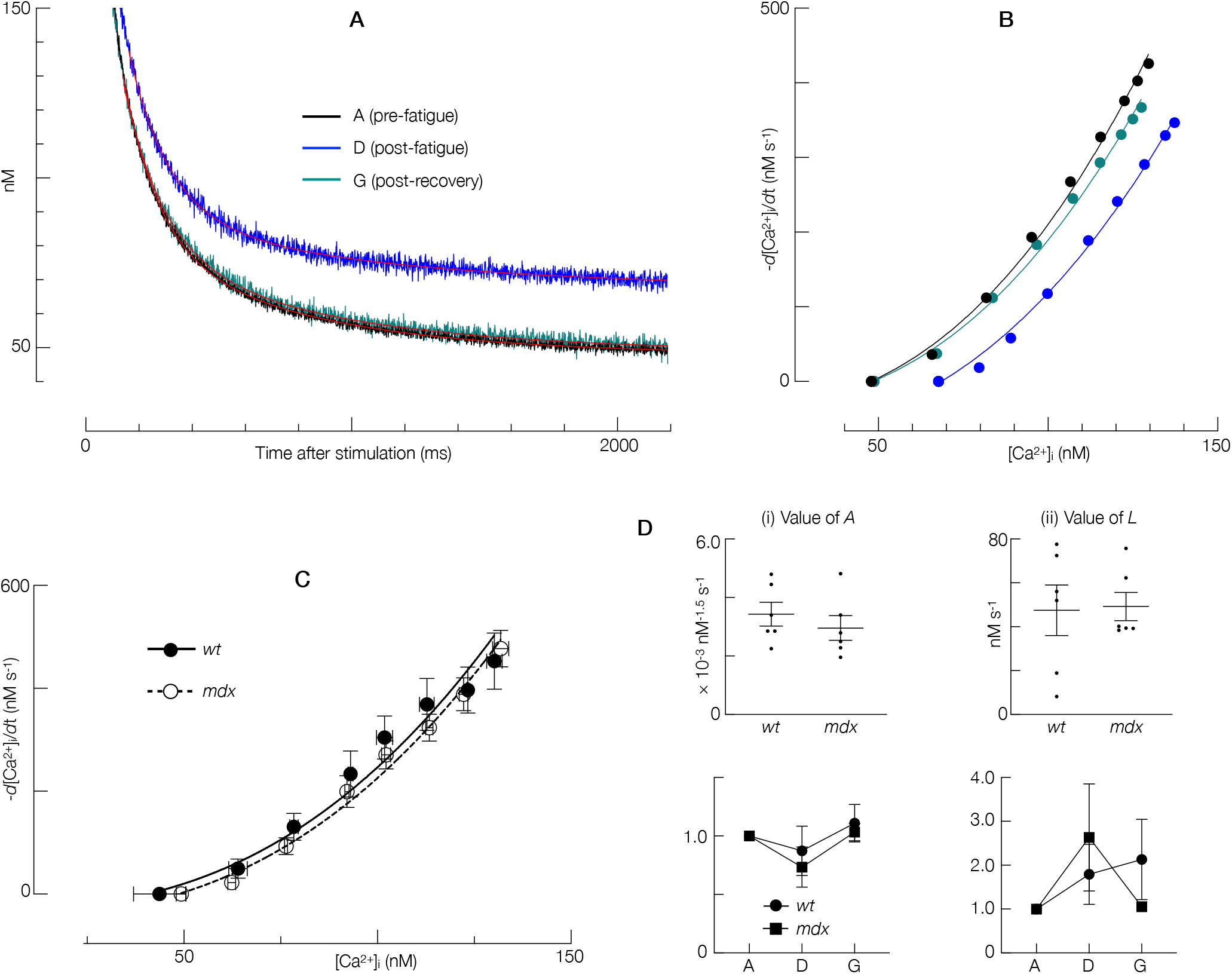
SR Pump function. **(A)** Slow decay phase of uncorrected [Ca^2+^]_i_ in three twitches from Stages A, D and G of an experiment in an *mdx* animal aged 7 wks. *Red lines* fitted double exponential functions. **(B)** SR pump function curves for these same twitches, showing the rate of [Ca^2+^]_i_ decline at different levels of [Ca^2+^]_i_. **(C)** SR pump function curves for all *wt* and *mdx* fibres at Stage A (before fatigue). The two curves are very similar. **(D)** Summary statistics for (i) the value of *A*; and (ii) the value of *L*. *Upper row* pre-fatigue results from Stage A. *Lower row* effects of fatiguing stimulation and recovery. Labels on *x*-axis refer to the stages of the experiment described in Figure 1. Values on *y*-axis are relative values, with Stage A given a value of 1.0.

These curves are called SR pump function curves and reflect the steady-state relationship between the concentration of myoplasmic Ca^2+^ and the rate of its removal by the SR.

It can be seen from Figure 3B that in this particular experiment, fatigue is associated with a reduced rate of Ca^2+^ removal at any level of myoplasmic [Ca^2+^], but after recovery this rate returns to pre-fatigue values.

#### Aggregate pump function curves

In Figure 3C, the points obtained in Figure 3B for the pre-fatigue condition were aggregated across all fibres and a single pump function curve fitted for each genotype. It is obvious that the curves for *wt* littermate controls and *mdx* are very similar, and there was no statistical difference between these two curves.

#### Pump function parameters

SR pump function can be characterised by the parameters *A* and *L* from Equation 1. *A* reflects pumping rate and *L* reflects leak rate. The parameter *N* must be set to a constant if different fibres and experimental stages are to be compared. With *N* set to 2.5, the value of *R*^2^ for the least-squares regression was above 0.97 for all fitted curves and above 0.99 for 90% of fitted curves. Figure 3D summarises the values of *A* and *L* across all fibres; the *upper row* shows pre-fatigue values and the *lower row* shows the values post-fatigue and post-recovery, relative to pre-fatigue. There was no difference in *A* or *L* between *wt* littermate controls and *mdx* before fatigue; the effects of fatigue and recovery were also no different between the two genotypes.

### [Ca^2+^]_i_ changes during tetanus before fatigue

Figure 4A compares the uncorrected [Ca^2+^]_i_ (*black line*) with the kinetically corrected [Ca^2+^]_i_ (*red line*) under 50-Hz stimulation in a fibre from a *wt* littermate control before fatigue. Note that the first spike of the kinetically corrected [Ca^2+^]_i_, produced by the first action potential of the stimulation train, is the largest. This is a characteristic feature of type MT-II fast [Ca^2+^]_i_ profiles (Calderón *et al.* 2009), and was observed in every kinetically corrected tetanus in this study.

**Figure 4.**
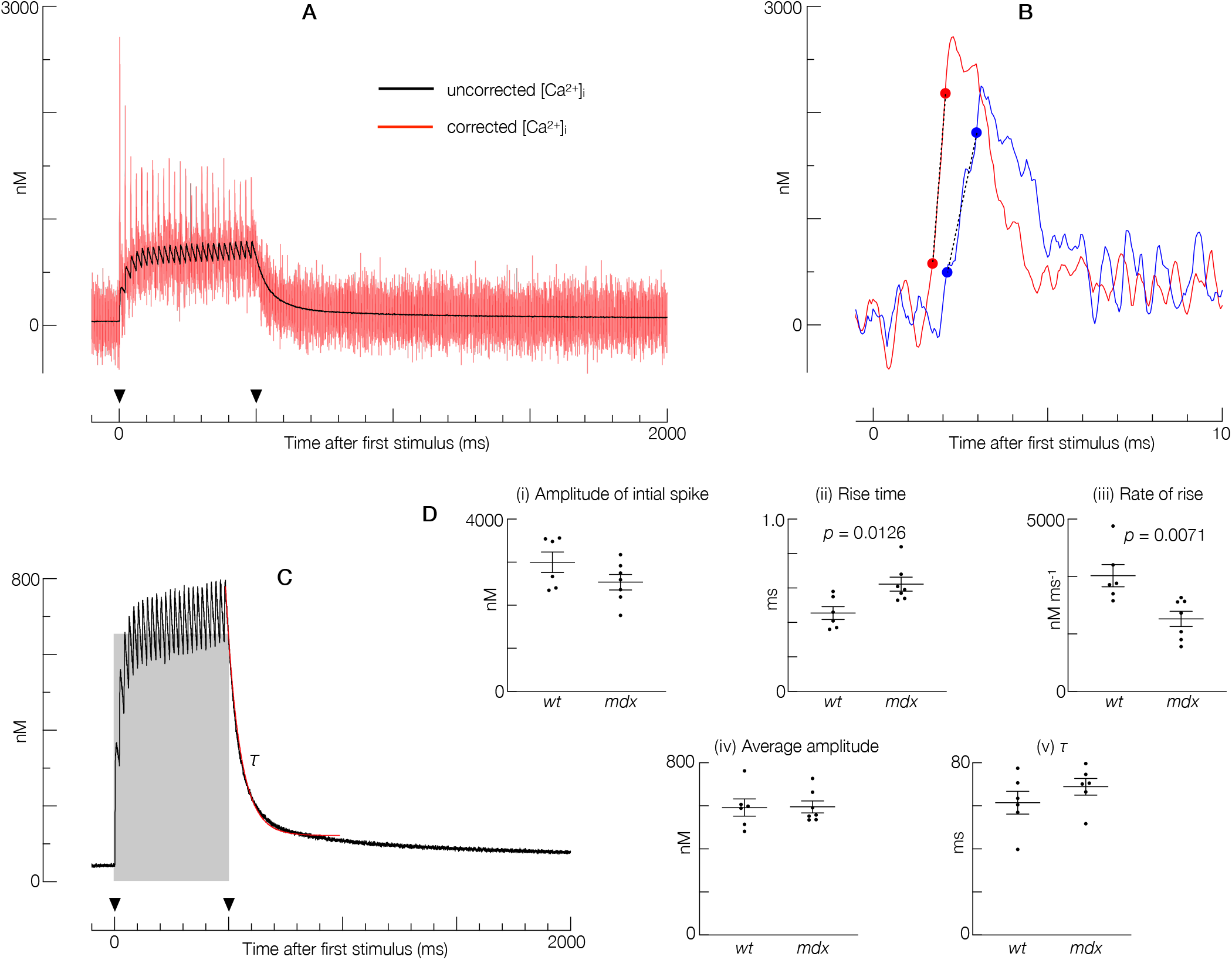
Changes in [Ca^2+^]_i_ during a tetanus. **(A)** Effect of kinetic correction of [Ca^2+^]_i_ in a *wt* littermate animal aged 7 wks. *Black line* uncorrected [Ca^2+^]_i_ at 50-Hz stimulation. *Red line* kinetically corrected [Ca^2+^]_i_. It can be seen that the kinetically corrected [Ca^2+^]_i_ consists of a large initial spike from the first action potential, followed by smaller spikes. *Arrowheads* indicate the start and end of stimulation. **(B)** *Red line* the first 10 ms of the corrected [Ca^2+^]_i_ shown in (A), focusing on the initial spike. *Blue line* kinetically corrected [Ca^2+^]_i_ from an *mdx* fibre, for comparison. *Dotted lines* join the points at which [Ca^2+^]_i_ reaches 20% and 80% of the amplitude. **(C)** *Black line* same uncorrected [Ca^2+^]_i_ as in (A). *Shaded rectangle* represents the integral of the uncorrected [Ca^2+^]_i_. *Curved red line* single exponential function with time constant *τ* fitted to the first 500 ms following the end of stimulation. **(D)** Summary statistics for (i) amplitude of the initial spike; (ii) rise time of the initial spike (timespan of the dotted line in (B)); (iii) rate of rise of the initial spike (slope of the dotted line in (B)); (iv) average amplitude of the tetanus; and (v) *τ*.

#### Amplitude of initial spike

The amplitude of the initial spike in Figure 4A is a sensitive indicator of Ca^2+^ release (Calderón *et al.* 2011). The amplitude of the initial spike in all fibres before fatigue is summarised in Figure 4Di. There was no significant difference between *wt* littermate controls and *mdx*.

#### Rise of initial spike

Figure 4B shows the large initial spike from Figure 4A at a higher time resolution. The kinetically corrected [Ca^2+^]_i_ from an *mdx* animal (*blue*) is also shown. The *dotted lines* join the points at which 20% and 80% of spike amplitude are reached. Figure 4Dii shows the time taken to rise from 20% to 80% of spike amplitude before fatigue in all fibres. This time was significantly longer in *mdx* than *wt* littermate controls (*p* = 0.0126). The rate of this rise (Figure 4Diii) was significantly slower in *mdx* than in *wt* littermate controls (*p* = 0.0071).

#### Average amplitude of tetanic [Ca^2+^]_i_

Figure 4C shows again the uncorrected [Ca^2+^]_i_ from the *wt* animal. The area of the *shaded rectangle* equals the integral of the [Ca^2+^]_i_ values from the start to the end of stimulation; its height therefore represents the average [Ca^2+^]_i_ throughout the period of stimulation. It was found that the integral of the uncorrected [Ca^2+^]_i_ and the integral of the kinetically corrected [Ca^2+^]_i_ were almost identical. Across all fibres before fatigue, the average amplitude of the tetanic [Ca^2+^]_i_ (Figure 4Div) was not significantly different between *wt* littermate controls and *mdx*.

#### Time constant of decay

In Figure 4C, the *red line* is a single exponential function with time constant *τ* fitted to the first 500 ms following the end of stimulation. Only the first 500 ms was fitted because during the fatigue runs there was only 0.5 s gap between stimulation trains. Across all fibres before fatigue, the time constant of decay (Figure 4Dv) did not differ between *wt* littermate controls and *mdx*.

### Effect of fatigue on tetanic [Ca^2+^]_i_

#### Average amplitude of tetanic [Ca^2+^]_i_

Figure 5A shows the uncorrected [Ca^2+^]_i_ of tetani at 12-s intervals during the fatigue run in a fibre from an *mdx* animal. The [Ca^2+^]_i_ at the start of each tetanus has been subtracted from the absolute [Ca^2+^]_i_ values so that all tracings start from the zero level; the left vertical axis thus records the [Ca^2+^]_i_ amplitude. The right vertical axis shows the average amplitude of [Ca^2+^]_i_ over the duration of each tetanus. Figure 5Div summarises the average amplitude at these six time points during the fatigue run in all fibres, as well as at post-fatigue (Stage E) and post-recovery (Stage H). Values are expressed relative to pre-fatigue values. There is a general fall in average amplitude during the fatigue run. Between the end of fatiguing stimulation and the tetanus measured at Stage E, there is virtually complete recovery to pre-fatigue levels, with little further change following the 3-min recovery period. However, there were no significant differences between *wt* littermate controls and *mdx* in these patterns.

**Figure 5.**
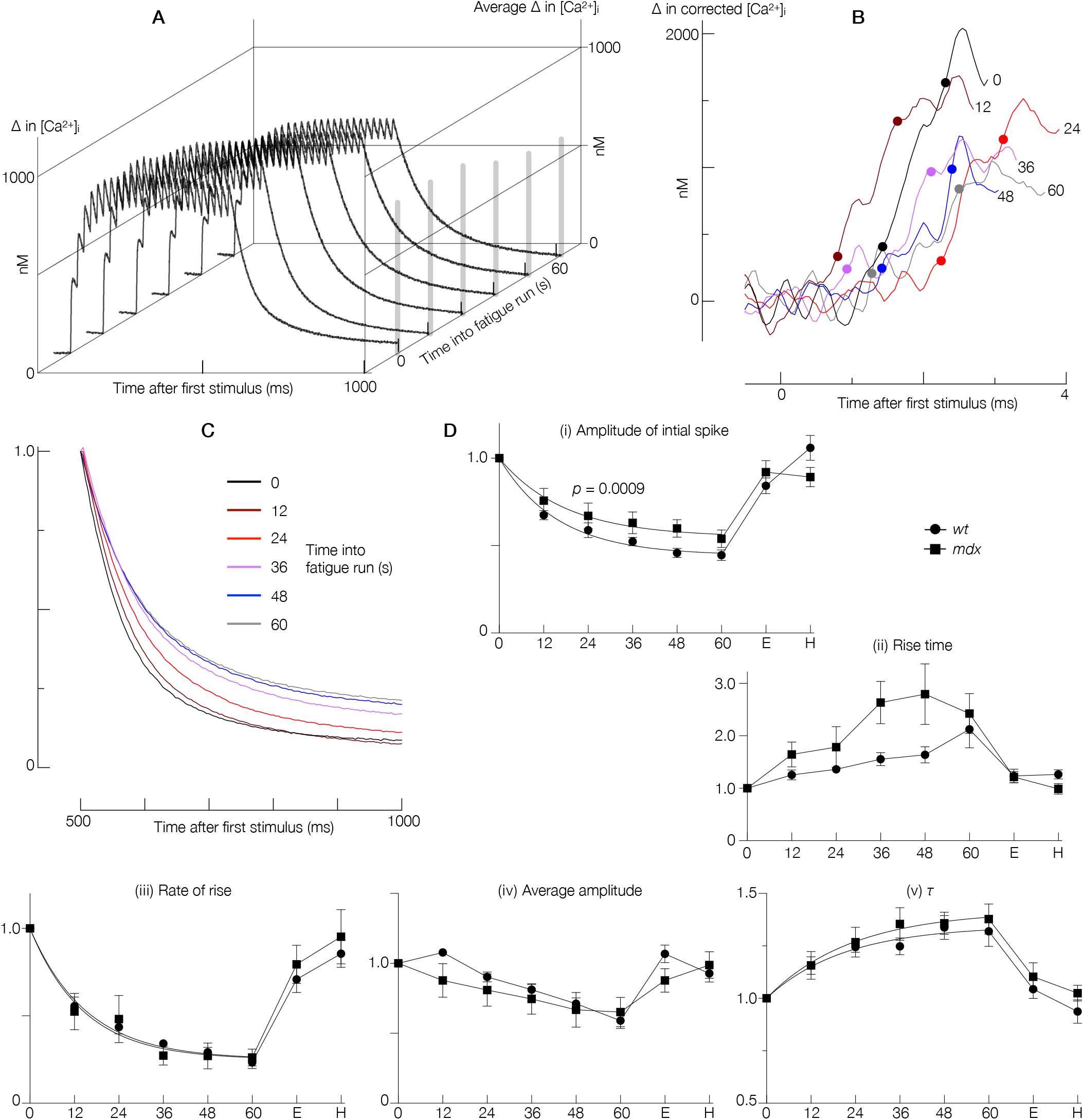
Effect of fatigue on the tetanic [Ca^2+^]_i_. **(A)** Uncorrected [Ca^2+^]_i_ for tetani at 12-s intervals during the fatigue run. *Right vertical axis* shows the average [Ca^2+^]_i_ amplitude of each tetanus over the period of stimulation. **(B)** The first few ms of the tetani in (A) after kinetic correction, showing the rising phase of the initial spike. The number on each curve indicates the time at which the tetanus occurred. The pair of dots on each curve indicate the times at which 20% and 80% of the spike amplitude are reached. **(C)** The decay phase of each tetanus in (A), with values normalised to the [Ca^2+^]_i_ at the end of stimulation. Data in (A), (B) and (C) taken from *mdx* animal aged 7 wks. **(D)** Summary statistics for the same parameters as in Figure 4(D), shown at 12-s intervals during the fatigue run and also at Stages E (post-fatigue) and H (post-recovery) of each experiment. All values have been normalised to pre-fatigue levels. In the cases of (i), (iii) and (v), it was possible to fit a single exponential function to the intra-fatigue data.

#### Amplitude of initial spike

Figure 5B shows the rising phase of the initial spike of kinetically corrected [Ca^2+^]_i_ for each tetanus shown in Figure 5A. Figure 5Di summarises how the amplitude of this initial spike changes across all fibres during the course of the experiment. For both *wt* littermate control and *mdx* fibres, it was possible to fit a single exponential function to the falling amplitude values during the fatigue run. The exponential curves for *wt* and *mdx* were significantly different (*p* = 0.0009). The diagram shows that the fall in amplitude of the initial spike during the fatigue run was less pronounced in *mdx* than in *wt*. At post-fatigue and post-recovery time points however, there were no significant differences, with *wt* and *mdx* reaching almost complete recovery in this parameter by Stage E of the experiment.

#### Rise of initial spike

On Figure 5B, the *dots* on each curve indicate the times at which 20% and 80% of the spike amplitude are reached. Figure 5Dii summarises how the 20%-80% rise time changes throughout the experiment in all fibres, while Figure 5Diii summarises the changes in the rate of rise. Figure 5Dii suggests that over the course of the fatigue run, the rise time is prolonged to a greater extent in *mdx* than in *wt* littermate controls. However, Figure 5Diii shows that the rate of rise over the course of the fatigue run is reduced to a similar extent in *wt* and *mdx*. The slowing of the rate of rise as fatigue progresses is well described by a single exponential decay function. The fitted exponential curves for *wt* and *mdx* are almost identical. The reason why the fatigue-associated reduction in rise rate is so similar in *wt* and *mdx*, when the fatigue-associated increase in rise time appears to be greater in *mdx*, is that the fatigue-associated reduction in spike amplitude is less in *mdx* (Figure 5Di). Following the fatigue run, both rise time and rise rate recovered to pre-fatigue levels.

#### Time constant of decay

Figure 5C shows the decay phase of the same uncorrected [Ca^2+^]_i_ as in Figure 5A, with values normalised to the [Ca^2+^]_i_ at cessation of stimulation to facilitate comparison. Figure 5Dv summarises the changes in the time constant of decay *τ* over the course of the experiment in all fibres. There was a fatigue-associated increase in *τ* that was well described by a single exponential function; however the fitted exponential curves for *wt* littermate controls and *mdx* were not significantly different. After the fatigue run, *τ* recovered to pre-fatigue levels in both *wt* and *mdx*.

## Discussion

We have examined the Ca^2+^ handling of the SR in unbranched fast-twitch muscle fibres from *mdx* mice aged 7 weeks. As described in the Introduction, we selected these fibres for two reasons. Firstly, branching in itself can affect the SR Ca^2+^ handling in dystrophin-deficient fibres, and we wanted to exclude this factor from our Ca^2+^ measurements. Secondly, fibres at this age have just regenerated after the period of massive necrosis, and would not have had sufficient time to accumulate other pathologies which could in themselves affect SR Ca^2+^ handling. (For the purposes of this discussion, we shall refer to fibres at this age as “immediately post-necrotic”, though it must be understood that the degenerative-regenerative cycle will continue thereafter at a stable low rate.) In addition, we have used wild-type littermates as control animals, thus avoiding the possibility that other genetic differences between *wt* and *mdx* mice could be affecting the SR Ca^2+^ handling.

Our fibre selection therefore allows us to move closer to the initial events resulting from a loss of dystrophin. Our findings indicate that the initial effects of dystrophin deficiency on SR Ca^2+^ handling are very slight. In our study, dystrophin deficiency did not affect the amplitude, rate of rise or decay time constant of the [Ca^2+^]_i_ response to a single depolarising pulse before fatigue. The effects of fatiguing stimulation and recovery on these parameters were also not different in the absence of dystrophin (Figure 2). SR pump function and SR leak in unfatigued fibres was not affected by dystrophin deficiency, and no effects could be unmasked after fatigue or after recovery (Figure 3). For tetani before fatigue, loss of dystrophin did not alter the amplitude of the initial spike, the average amplitude or the decay time constant of [Ca^2+^]_i_ (Figure 4). The time course of fatigue-induced changes in the rise time of the initial spike, rate of rise of the initial spike, average amplitude and decay time constant were also unaffected by dystrophin deficiency (Figure 5).

As muscular dystrophy in the *mdx* mouse is widely regarded as a disease of pathological Ca^2+^ handling (Burr & Molkentin, 2015), it may seem surprising that our present study found so little difference between the [Ca^2+^]_i_ responses of *wt* littermate control and *mdx* muscle fibres to electrical stimulation. However, it must be remembered that we used unbranched, immediately post-necrotic *mdx* fibres, so we were observing early events following the loss of dystrophin. As time progresses, the fibres degenerate and branch, and this degeneration and branching would in themselves magnify the Ca^2+^ pathology. It has traditionally been thought that loss of dystrophin would de-stabilise the sarcolemma, leading to contraction-induced tears and an influx of Ca^2+^ (Petrof *et al.* 1993; Hopf *et al.* 1996). However, the largely normal Ca^2+^ handling of our study fibres suggests that the early mechanisms of Ca^2+^ pathology in dystrophin deficiency may be more subtle, such as an excessive ROS-mediated activation of stretch-activated Ca^2+^ channels in the sarcolemma (Allen *et al.* 2016).

The two differences that we did find relate to SR Ca^2+^ release. Firstly, the initial spike in [Ca^2+^]_i_ during a tetanus had a significantly longer rise time and slower rate of rise in the absence of dystrophin (Figure 4D, ii and iii). The kinetics of this initial spike are a sensitive indicator of SR Ca^2+^ release function (Calderón *et al.* 2011). An impairment of Ca^2+^ release is one of the more consistent findings in SR Ca^2+^ studies on *mdx* mice (Plant & Lynch 2003; Woods *et al.* 2004; Hollingworth *et al.* 2008), so our results suggest that this may be one of the earliest manifestations of Ca^2+^ pathology in the SR of *mdx* mouse skeletal muscle.

Secondly, the exponential decline in the amplitude of this initial spike as fatigue progresses was ameliorated in the absence of dystrophin (Figure 5D, i). One explanation for this may lie in the fact that fatigue impairs Ca^2+^ resequestration by the SR (Stephenson *et al.* 1998), but as the SR in *mdx* muscle may have a chronically reduced [Ca^2+^] (Robin *et al.* 2012), the SERCA pump in *mdx* muscle would work against a smaller gradient and the impairment of re-uptake would be less pronounced, mitigating the fall in the amount of releasable Ca^2+^.

It has been reported that dystrophin deficiency renders the SR especially leaky to Ca^2+^, both in mechanically stressed fibres (Wang *et al.* 2005; Bellinger *et al.* 2009) and in unstressed fibres (Divet & Huchet-Cadiou 2002, Robin *et al.* 2012). Our SR pump function analysis detected no such changes (Figure 3). None of the aforementioned studies explicitly excluded branched fibres. Divet & Huchet-Cadiou (2002) and Bellinger *et al.* (2009) used EDL muscle, which contains a higher proportion of branched fibres than FDB muscle. Robin *et al.* (2012) used FDB fibres from mice aged 1-2 months, so it is possible that they included some fibres which were still in the peak necrotic stage. Only Wang *et al.* (2005) used FDB fibres of a comparable age to those in our study, but they subjected their fibres to osmotic shock. Lovering *et al.* (2009) used a similar hyper-osmotic challenge, but explicitly distinguished between branched and unbranched fibres. They found that unbranched *mdx* fibres had significantly greater Ca^2+^ spark activity than *wt* fibres. Hence mechanical stress may be required to unmask a greater leakiness of the SR in dystrophin deficiency.

To summarise, we have found that the SR Ca^2+^ handling characteristics of unbranched, immediately post-necrotic fast-twitch fibres from *mdx* mice are largely similar to those of fibres from *wt* littermates. This suggests that the early effects of dystrophin deficiency on SR Ca^2+^ handling are very subtle, limited perhaps to an impairment of Ca^2+^ release.

